# Structural variants at the *BRCA1/2* loci are a common source of homologous repair deficiency in high grade serous ovarian carcinoma

**DOI:** 10.1101/2020.05.11.088278

**Authors:** Ailith Ewing, Alison Meynert, Michael Churchman, Graeme R. Grimes, Robert L. Hollis, C. Simon Herrington, Tzyvia Rye, Clare Bartos, Ian Croy, Michelle Ferguson, Mairi Lennie, Trevor McGoldrick, Neil McPhail, Nadeem Siddiqui, Suzanne Dowson, Rosalind Glasspool, Melanie Mackean, Fiona Nussey, Brian McDade, The Scottish Genomes Partnership, Lynn McMahon, Athena Matakidou, Brian Dougherty, Ruth March, J. Carl Barrett, Iain A. McNeish, Andrew V. Biankin, Patricia Roxburgh, Charlie Gourley, Colin A. Semple

## Abstract

Around half of high grade serous ovarian carcinomas (HGSOC) show homologous recombination repair deficiency (HRD), often caused by germline or somatic single nucleotide variant (SNV) mutations or small indels disrupting *BRCA1/2*. We have uniformly processed the largest collection of whole genome sequencing (WGS) data from HGSOC samples to date (N=205), comprehensively characterising the somatic mutational landscape, and expression at the *BRCA1/2* loci. We discover that large structural variants (SV) are a frequent but unappreciated source of *BRCA1/2* disruption in these tumours. Somatic structural variation at these loci is dominated by multi-megabase deletions that span the entirety of *BRCA1* (median = 4.9Mb) or *BRCA2* (median = 6.2Mb), independently affecting a substantial proportion of patients (16%) in addition to those affected by damaging germline or somatic short variants, within the *BRCA1/2* coding sequences (24%). In common with previous studies, we show that the presence of damaging somatic SNVs or short indels in *BRCA1* (OR=10, 95% CI 1.8-103, p=0.002, adj p=0.027 and *BRCA2* (OR=17, 95% CI 2.1-816), p=0.002, adj.p=0.021) was found to influence HRD. For the first time we also study the compound effect of SV and SNV or short indel mutations at both loci, demonstrating that SVs often contribute to compound deficiencies involving SNVs or indels, with large somatic deletions contributing to these compound deficiencies in 15/205 (7%) of samples. Notably the strongest risk of HRD (OR=19 (2.4-896), p=6.6×10^-3^, adj P=8.5×10^-3^) is generated by combined large deletions at *BRCA1* and *BRCA2* in the absence of SNVs or indels, affecting 3% of patients. Overall, we show that HRD is a complex phenotype in HGSOC tumours, affected by the patterns of shorter variants such as SNVs and indels, SVs, methylation and expression seen at multiple loci, and we construct a successful (ROC AUC = 0.75) predictive model of HRD using such variables. In addition, HRD impacts patient survival when conferred by mechanisms other than through the well-understood short variants at *BRCA1/2*, currently exploited in the clinic. These results alter our understanding of the mutational landscape at the *BRCA1/2* loci in highly rearranged tumours, and increase the number of patients predicted to benefit from therapies exploiting HRD in tumours such as PARP inhibition.

## Main

Homologous recombination repair deficiency (HRD) is identifiable in many cancers and is particularly prominent in high grade serous ovarian cancer (HGSOC)^1^, affecting around 50% of tumours^2^ and leaving detectable mutational spectra across the tumour genome^3^. The mutational landscape of HGSOC is dominated by extensive genomic copy number changes and structural rearrangement driven by chromosome instability and defective DNA repair, rather than the patterns of recurrent point mutation in tumour suppressor and oncogenes often observed in other solid tumours^4,5^.

Germline variants disrupting the coding sequence of *BRCA1* and *BRCA2* are the most common types of HRD-associated defect, occurring in 8% and 6% of patients respectively, while disruptive somatic short mutations in these genes are present in an additional 4% and 3% of patients respectively^6,7^. These germline short variants (GSV) and somatic short variants (SSV) include single nucleotide variants (SNVs) as well as short indels, with frameshifts being the predominant mechanism of inactivation. These *BRCA*-deficient patients represent approximately 20% of patients with HGSOC. An additional 11% of patients are thought to be *BRCA*-deficient through epigenetic silencing of *BRCA1*^2,8^. Mutational or epigenetic inactivation of other genes involved in the HR pathway are also thought to confer HRD in a smaller proportion of HGSOC patients^7,9–12^. Genome-wide patterns of SNVs, indels and structural variation have been identified as strong predictors of *BRCA1/2* deficiency^3^. These mutational signatures of *BRCA1/2* deficiency are also found in additional patients who lack short variants at *BRCA1/2*, suggesting that other unknown aberrations may also be involved in HRD^3^. The demonstration of *BRCA1/2* loss and detection of HRD is crucial in the management of HGSOC and other cancers to identify patients whose prognosis is markedly improved by the administration of PARP inhibitors^13–15^. PARP inhibitors selectively kill cells that are deficient in HR (homology-directed repair) because these cells can neither resolve stalled replication forks nor accurately repair the increased number of double strand breaks that result from the use of these agents^16^.

The clinical importance of germline and somatic short variants at *BRCA1/2* is well established in cancer, with variants documented in multiple repositories^17,18^. In contrast, the abundance and effects of structural variants (SVs) at *BRCA1/2* are not well understood, particularly for large SVs encompassing multi-megabase regions. Similarly, the compound effects of SVs and short variants occurring simultaneously at *BRCA1* and *BRCA2* are poorly studied. Matched tumour-normal whole genome sequencing (WGS) of freshly-frozen tissue is accepted as the best resource to accurately detect SVs in tumours but in the past such data have been scarce for HGSOC^19,20^. Here we comprehensively characterise the mutational landscape of *BRCA1/2* in HGSOC using the largest collection to date of uniformly processed WGS data (N=205), comprising two previously published cohorts^5,6^, as well as a large novel cohort described here for the first time. We document the prevalence of HRD across these three cohorts to reveal the complexity of the mutations associated with HRD, their impact on gene expression and associations with clinical variables.

## Results

WGS data from matched primary tumour and normal blood samples were uniformly remapped and analysed to generate a range of somatic mutation calls (Methods, Supplementary Figure 1) for three HGSOC cohorts: the Australian Ovarian Cancer Study^5^ (AOCS) (N=80), The Cancer Genome Atlas (TCGA)^6^ WGS HGSOC samples (N=44) and the previously unpublished Scottish High Grade Serous Ovarian Cancer (SHGSOC) study (N=81). The combined uniformly analysed cohort (N=205) presented here represents the largest collection of HGSOC WGS data investigated to date.

### Large structural variants are a frequent source of *BRCA1/2* disruption in HGSOC

We identified SVs in HGSOC samples using a combination of three algorithms chosen in order to enable the most accurate detection of the full range of SVs in the tumours. All large (>1Mb) deletion and duplication calls were identified by two variant callers that incorporated two independent forms of evidence: read depth variation and deviations in heterozygous single nucleotide variant (SNV) allele frequencies. All calls that passed manual curation were then carried forward for analysis (Supplementary Table 2). During the course of these analyses, the WGS data from the AOCS and TCGA were also reprocessed and analysed in parallel by the Pan-Cancer Analysis of Whole Genomes (PCAWG) project^21^. Recent studies have suggested that ensemble approaches to SV detection are overly conservative^22,23^ but we found that our manually curated CNA calls >1Mb demonstrated a high correspondence with PCAWG variant calls. Of the large copy number variants identified using our approach in samples included in PCAWG, 98% (41/42) were also recovered independently by the PCAWG project within the same samples, suggesting that our SV calls are likely to carry a negligible false positive rate.

A variety of SVs were detected at the *BRCA1/2* loci but were dominated by large multimegabase deletions spanning the entirety of *BRCA1* or *BRCA2*. In all three cohorts these deletions often encompass a large proportion of chromosomes 17 or 13 though the majority are more focal (median *BRCA1* deletion = 4.9Mb, median *BRCA2* deletion = 6.2Mb) (Figure 1). Heterozygous deletions occur at similar rates at *BRCA1* (16%) and *BRCA2* (14%) overall, and at comparable rates between the similarly sized AOCS and SHGSOC cohorts (Supplementary Table 1). In 6/205 (3%) of samples in the combined cohort, we observe large deletions at both *BRCA1* and *BRCA2* in the absence of *BRCA1/2* short variants. This is in contrast to the mutually exclusive pattern of mutation observed for short variants at *BRCA1/2* where a sample only ever has at most one short variant across *BRCA1* and *BRCA2*. Inversions occur less often than deletions, but do occur in isolation in 6% of samples, and within groups of large overlapping inversions in 5% of samples. In addition, we observe large duplications that span the entire length of either gene in all cohorts (Supplementary Table 2). We observe similar genome-wide SV mutational spectra in each cohort despite the clinical differences among them: in particular AOCS represents chemoresistant/relapsed cases, while SHGSOC is composed mainly of samples taken before treatment. This suggests that large SVs predicted to impair *BRCA1/2* function are a general feature of HGSOC evolution and should be considered alongside *BRCA1/2* short variants when investigating the functional impact of mutational aberrations at *BRCA1/2*. Large deletions at *BRCA1/2* are abundant but are not significantly over-represented at these genes relative to the rates of large deletions throughout the genome in HRD samples (Supplementary Figure 2). However, given the critical roles of *BRCA1/2* in DNA repair, deletions at these loci may have a disproportionate effect on function.

**Figure 1:**
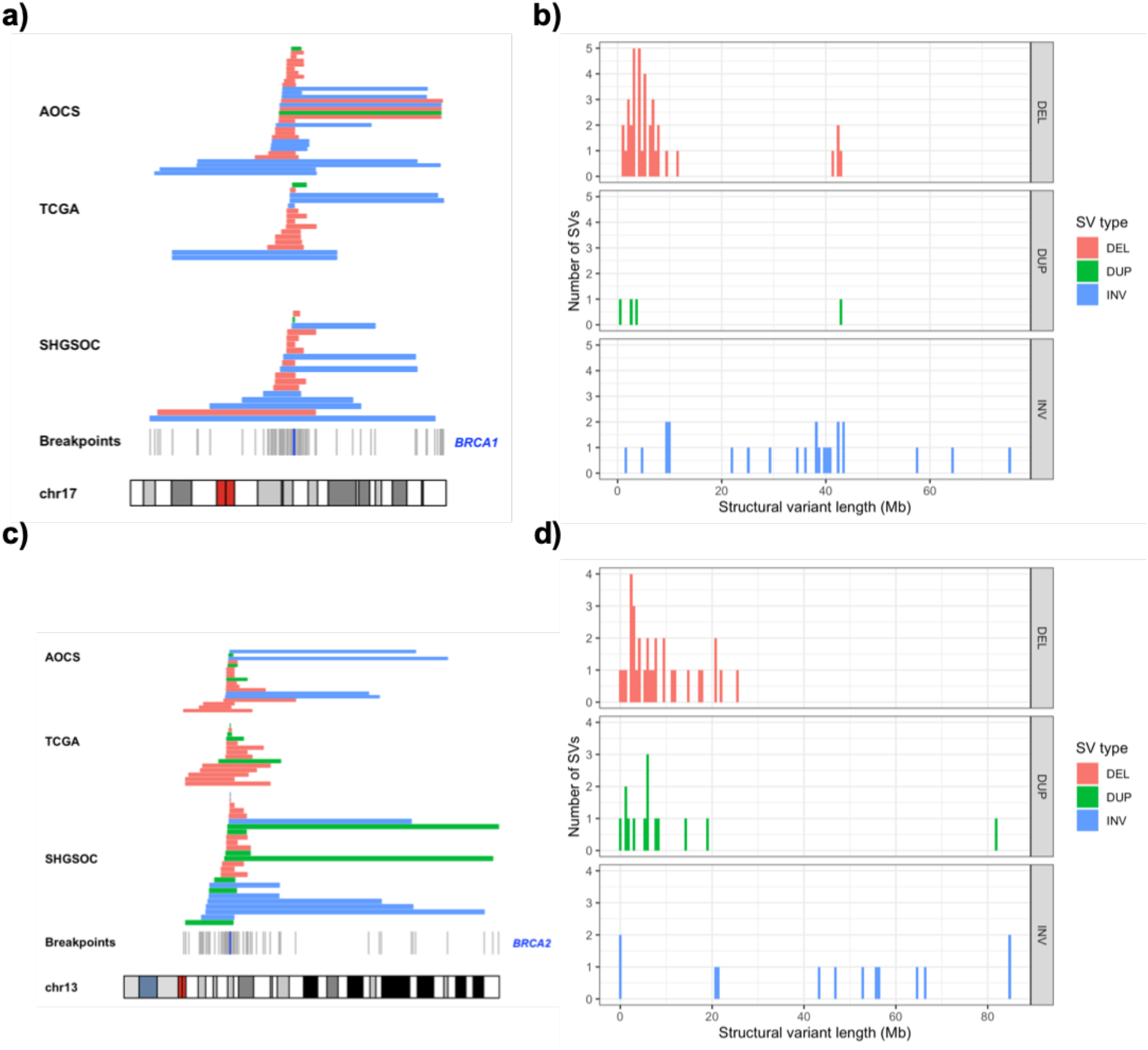
Abundance, location and size of structural variants overlapping *BRCA1/2* in three HGSOC cohorts. a) Alignment of structural variants overlapping *BRCA1* across the AOCS, TCGA and SHGSOC cohorts with breakpoints marked in grey according to their position on chromosome 17. Location of *BRCA1* marked by a blue line with deletions (red), duplications (green) and inversions (blue). b) The distribution of sizes of structural variants (Mb), overlapping *BRCA1* across all cohorts. c) Alignment of structural variants overlapping *BRCA2* across the three cohorts with breakpoints marked in grey according to their position on chromosome 13. Location of *BRCA2* marked by a blue line. d) The distribution of sizes of structural variants (Mb) overlapping *BRCA2* across all cohorts with deletions (red), duplications (green) and inversions (blue).

### Large deletions spanning *BRCA1/2* contribute to HRD independently of pathogenic SNVs and indels

We examined the functional impact of all *BRCA1/2* mutations detected across all cohorts using an established method, HRDetect^3^, which predicts HRD based upon genome-wide mutational spectra, and therefore provides a functional readout for the HR repair pathway in tumours (Figure 2).

**Figure 2:**
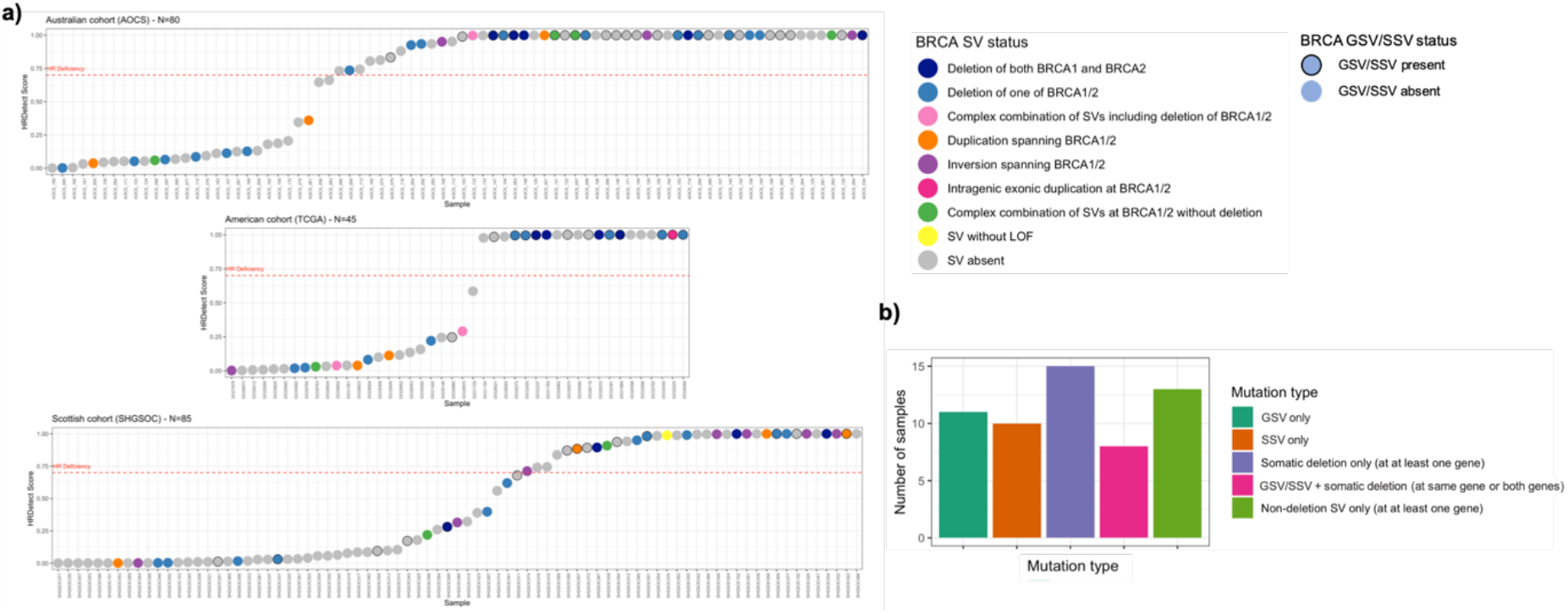
Homologous repair deficiency across three HGSOC cohorts. a) Predictions of HRD for three large cohorts of HGSOC coloured by *BRCA1/2* mutation status. Categories of mutation include GSVs and SSVs, 6 types of structural variation and absence of any *BRCA1/2* variant whether it be an SV or a short variant. The HRDetect scores range from 0, least likely to be HR deficient to 1, most likely to be HR deficient. The red dashed line represents the threshold of 0.7 representing HRD^3^. b) The number of HRD tumours with different categories of *BRCA1/2* short variants or deletions.

A high proportion (86%) of tumours with damaging GSVs or SSVs in *BRCA1/2* were assigned HRDetect scores indicating HRD (>0.7). As expected, patients with short variants in *BRCA1/2* even in the absence of deletion, are more likely to have HR deficient tumours than those patients without short or structural variants at these genes (GSV OR 6.9, 95% CI 1.8 – 33, p-value=2.3×10^-3^, adj. p-value= 3×10^-2^; SSV OR 25, 95% CI 3.2-1121, p-value = 1.7×10^-4^, adj. p-value = 2.3×10^-3^) (Figure 3). Four samples with GSVs at *BRCA1/2* demonstrated low HRDetect scores and accordingly showed no evidence for subsequent loss of the wild-type allele in the tumour which suggests that certain GSVs that are predicted to be disruptive are insufficient to generate HRD (Supplementary Table 3).

**Figure 3:**
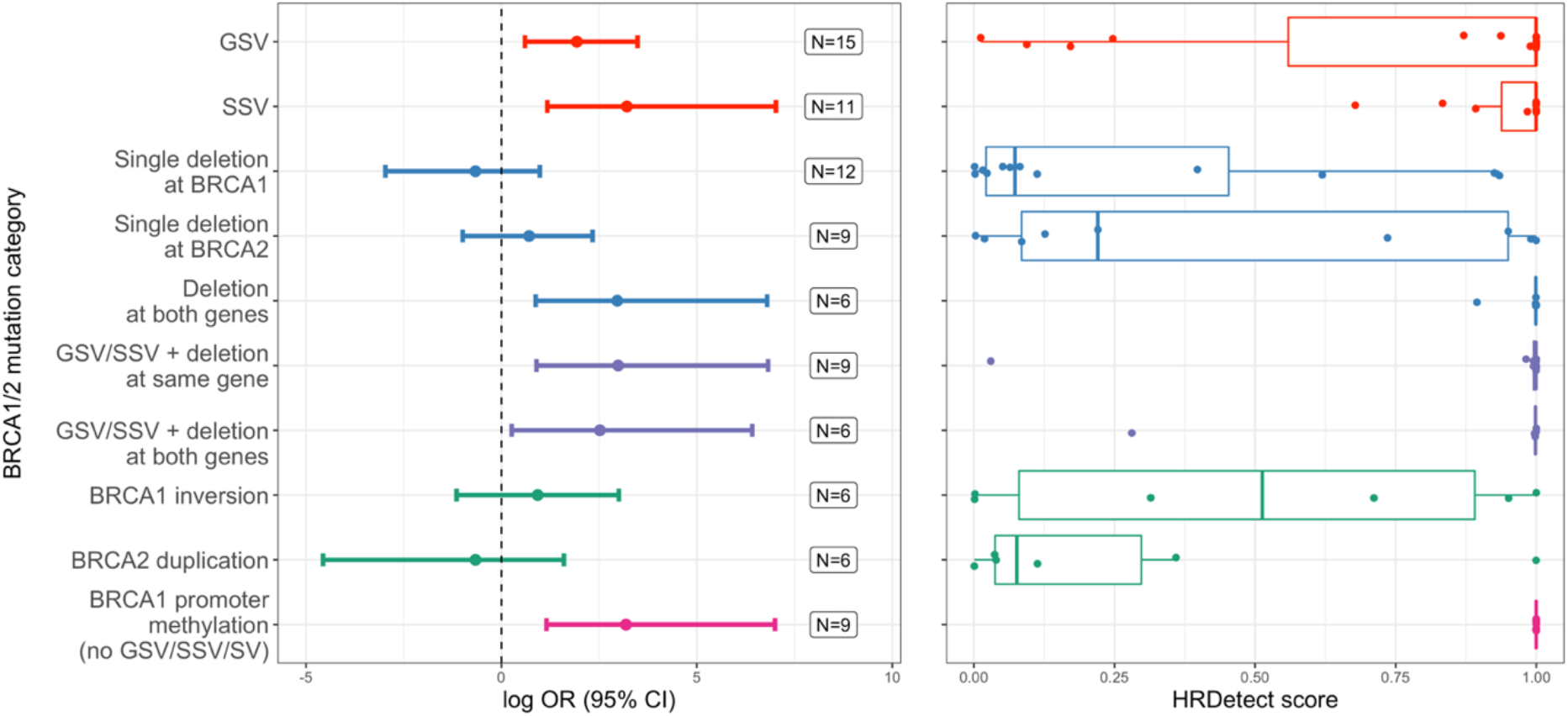
*BRCA1/2* mutation classes and repair deficiency in three HGSOC cohorts. The increase in log odds ratio of HRD (HRDetect score > 0.7) associated with different categories of mutation at *BRCA1/2* in comparison to the frequency of the reference category where samples lack any evidence of *BRCA1/2* inactivation (GSV, SSV, SV or methylation). ORs are defined using Fisher’s Exact tests for enrichment. Error bars represent 95% confidence intervals. Mutually exclusive categories of mutation examined include GSV only, SSV only, the presence of a deletion at one or both genes without a GSV or SSV, the presence of a short variant together with deletion of one or both genes, non-deleting SVs in samples without short variants or deletions, samples with *BRCA1* promoter methylation and no mutational *BRCA1/2* deficiencies. Samples with *BRCA1* promoter methylation are excluded from all *BRCA1/2* mutational categories.

The majority of samples (65%) with *BRCA1/2* deletions were assigned high HRDetect scores indicating HRD, though many occur in tumours that also harbour short variants. Many (30/49, 61%) of the samples with *BRCA1/2* deletions lack short variants, permitting the analysis of the effects of deletions independently of short variants. The compound effect of deletions at both the *BRCA1* and *BRCA2* loci in the absence of short variants is particularly pronounced, demonstrating a significantly increased risk of HRD (OR 19, CI 2.4-896, p=1.3×10^-3^, adj p =1.7×10^-2^) and consistently high HRDetect scores. In fact, compound deletions at both loci generate an OR that is comparable with other classes of HRD mutations known to have clinical importance, including the well-studied disruptive *BRCA1/2* short variants (Figure 3). Single deletions at either *BRCA1* or *BRCA2* do not consistently confer an increased risk of HRD in the absence of a *BRCA1/2* short variant (Figure 3) though the analysis may be underpowered to detect small effects given the current sample size. The HRDetect scores for samples with these single deletions form a bimodal distribution for which we have been unable to find a defining characteristic for the difference, such as the length of the deletion, resultant level of gene expression or a background of whole genome doubling. The estimated effect that we observe of a single deletion at *BRCA2* merits further investigation although it does not achieve statistical significance in our data (OR 2, 95% CI 0.37-10.3) (Figure 3). Given the notable effect of compound deletions in the absence of *BRCA1/2* short variants we conclude that large *BRCA1/2* deletions have a currently unexploited potential as biomarkers in patient stratification for treatments targeting HRD particularly when they occur together.

Beyond deletions the functional impact of other classes of SV, such as inversions or duplications, is less well studied but it is clear that samples bearing these mutational classes can show evidence of HRD (Figure 2). Inversions can occur at either gene, but are more likely at *BRCA1*, and duplications are more likely to occur at *BRCA2* (Supplementary Table 1). The 6 samples with only *BRCA1* inversions split equally into HRD and HR proficient groups, which suggests that in isolation their presence is not associated with HR deficiency. In contrast, only one of the samples with only *BRCA2* duplications is HR deficient, which suggests potential for enrichment in HR proficient samples but this would need to be further explored in greater sample sizes (Figure 3). The potential role of these sorts of events in repair deficiencies within tumours is intriguing particularly as it is less clear what the mechanism of action of these events might be.

### Deletions are a frequent source of biallelic *BRCA1/2* inactivation in repair deficiency

Samples across the combined cohort never had more than one short variant across *BRCA1* and *BRCA2*. This suggests that somatic short variants (SSVs) are not a mechanism for biallelic inactivation of a gene affected by a germline short variant (GSV) and also that deficiency caused by a short variant at one gene is mutually exclusive with deficiency achieved by a short variant at the other. This is consistent with reports from a previous study of HGSOC^24^ (Supplementary Table 3). In contrast, of the HGSOC tumours with a GSV at *BRCA1/2* predicted to cause HRD we find that 11/32 (34%) show evidence for an SV at the same gene which, in combination with the GSV, may contribute to HRD if the SV occurs on the other allele from the GSV (Supplementary Tables 2 and 3). In the combined HGSOC cohort we find that most of these somatic events (8/11 = 73%) are large deletions, while a further two tumours possess more than one SV spanning the same gene as the GSV, in the absence of a somatic deletion, and one more shows evidence of somatic duplication. The importance of ‘second hit’^25^ mutations in tumours is well established^26^ but these data suggest that multimegabase deletions have an under-appreciated role in this phenomenon in HGSOC. Across the three cohorts, 24% (50/205) of patients have a disruptive short variant at either *BRCA1* or *BRCA2*, and 30% (15/50) of these patients also carry a *BRCA1/2* deletion at the same locus. Also, it appears that SVs, including deletions, can occur at both *BRCA1* and *BRCA2* in the same sample. Large somatic deletions occur at both *BRCA1* and *BRCA2* in 13 samples and in 7 of these samples there is no short variant at either gene, although 1 sample had a hypermethylated *BRCA1* promoter. These data suggest that large deletions and other SVs disproportionately contribute to biallelic inactivation in HGSOC, driven by the unusually high rates of structural variation seen in this cancer^27^.

### Deletions spanning *BRCA1/2* are associated with lower gene expression

We found that patients with a deletion overlapping *BRCA1*, in the absence of a GSV or SSV, had lower *BRCA1* expression than those patients with no *BRCA1* short or structural variants (log2 fold change of no GSV/SSV/SVs versus a deletion= 0.45, p-value=0.0093). We also found that tumours with large somatic deletions at *BRCA2* had lower *BRCA2* expression than those without GSV/SSV/SVs at *BRCA2* (log2 fold change in expression between no GSV/SSV/SV versus a deletion = 0.43, p-value = 0.037) (Figure 4d). In spite of the unavoidable heterogeneity in tumour expression data among samples, the trends observed here are consistent with a direct effect of large *BRCA1/2* deletions, inducing HRD by reducing *BRCA1/2* expression, though indirect mechanisms cannot be excluded.

**Figure 4:**
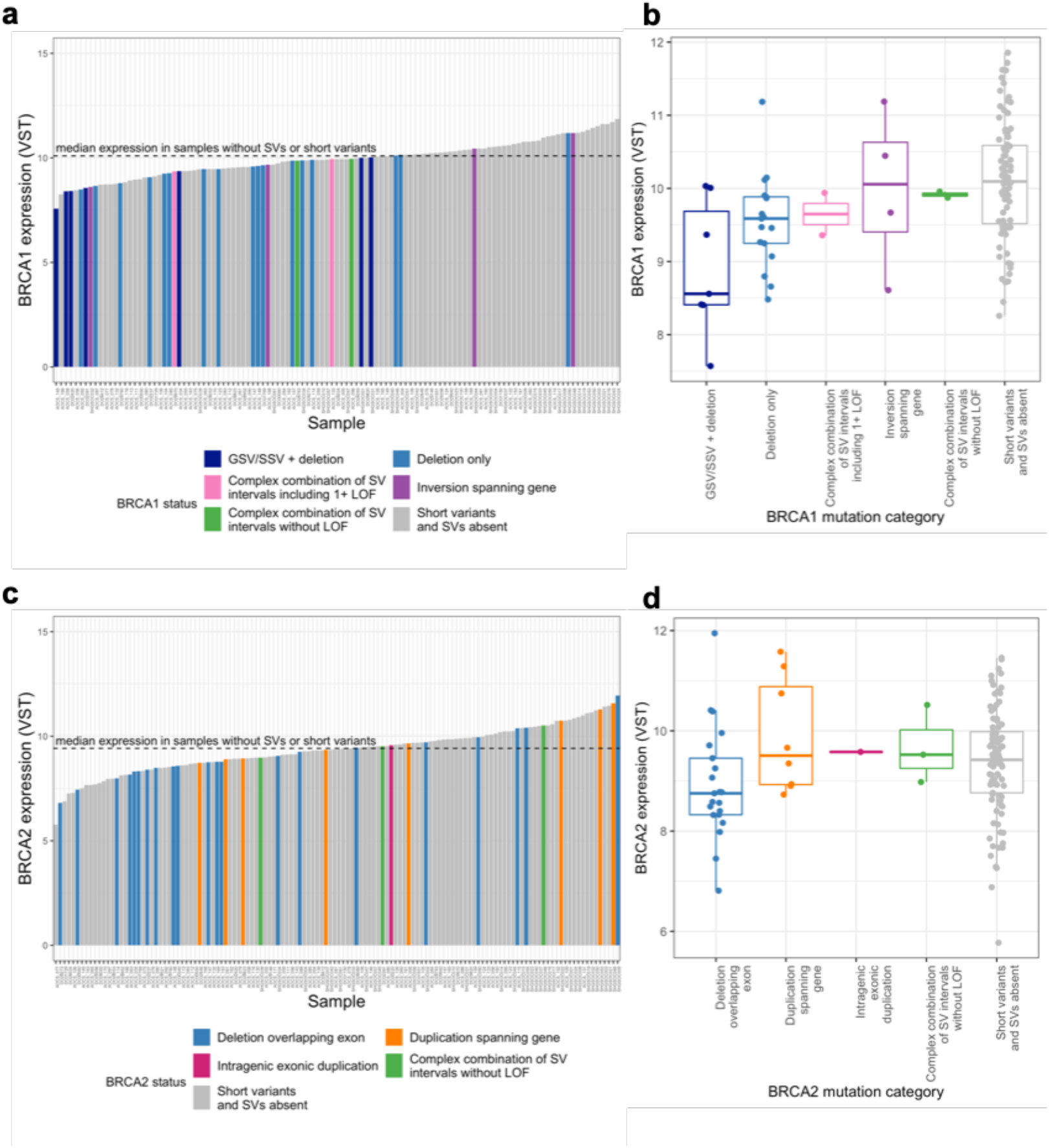
Structural variation and expression of *BRCA1/2* in three cohorts HGSOC. a)c) Expression of *BRCA1/2* (variance stabilising transformed RNA-seq counts) across samples ordered from lowest to highest expression. Median *BRCA1/2* expression is indicated by a black dashed line. Sample bars are coloured by *BRCA1/2* mutational category. b)d) Boxplot of *BRCA1/2* expression for each category of *BRCA1/2* mutation. The *BRCA1* deletion category is split into those samples with SNVs and deletions and those with only deletions as their expression is significantly different (Supplementary Figure 3a). This is not the case for *BRCA2* so all samples with deletions are considered together to maximise the available sample size (Supplementary Figure 3b).

Exploiting our novel combined cohort with matched genomic and transcriptomic data, we identified a list of differentially expressed (DE) genes between HR deficient and HR proficient HGSOC tumours, encompassing 306 protein coding genes (Supplementary Table 4). Notably these genes do not include known HR genes and given their diverse functions their dysregulation is likely to be a consequence rather than a cause of HRD. The variation in these genes, as defined by their first principal component, is significantly different between HR deficient and HR proficient samples in the combined cohort (Wilcox p-values =2.3×10^-10^) (Supplementary Figure 4a), 4b)). Transcriptomic signatures have previously been generated^28–31^ to identify HRD tumours; however, most have used suboptimal proxies such as mutation rate to predict HRD or have been based upon expression in HR deficient cell lines or samples that are not from HGSOC patients^28–31^. When we identified DE genes between HR deficient and HR proficient samples in a subset of the cohort the expression of these genes failed to accurately discriminate HR deficient from HR proficient samples in the unexamined remainder of the cohort (Wilcox p-value=0.92)(Supplementary Figure 4c), 4d)). This is consistent with previous reports^32^ and suggests that although the transcriptome is perturbed in the presence of HRD, such perturbations are not consistent, and consequently these expression changes are poor predictors of HRD.

### Integrative modelling reveals complex mechanisms underlying repair deficiency

We comprehensively modelled the effects of a range of genomic alterations at the *BRCA1/2* loci on HRD, to investigate the relative importance of these features in explaining the patterns of HRD observed. Given the relative sparsity of the data and the correlation between features we used a multivariable elastic net regularised regression model.

In addition to the previously reported impact of short variants at *BRCA1/2* and *BRCA1* promoter hypermethylation on HRD, large deletions at *BRCA2* confer an increased risk of HRD. Furthermore, samples with double deletions, where deletions are found at both *BRCA1* and *BRCA2*, are more likely to be HR deficient. Importantly, the influence of these double deletions on HRD exceeds that of genome-wide large CNV loads and genome-wide SV loads. Also, large inversions at *BRCA1* are independently associated with an increased risk of HRD. The functional impact of these events on the gene is currently unknown but this suggests that these events may either be markers for processes that impact the gene’s function or may even directly impact the function of the gene themselves. The model’s ability to predict HRD was good with a mean ROC curve AUC of 0.75, which although promising suggests that there are additional unknown sources of HRD (Figure 5).

**Figure 5:**
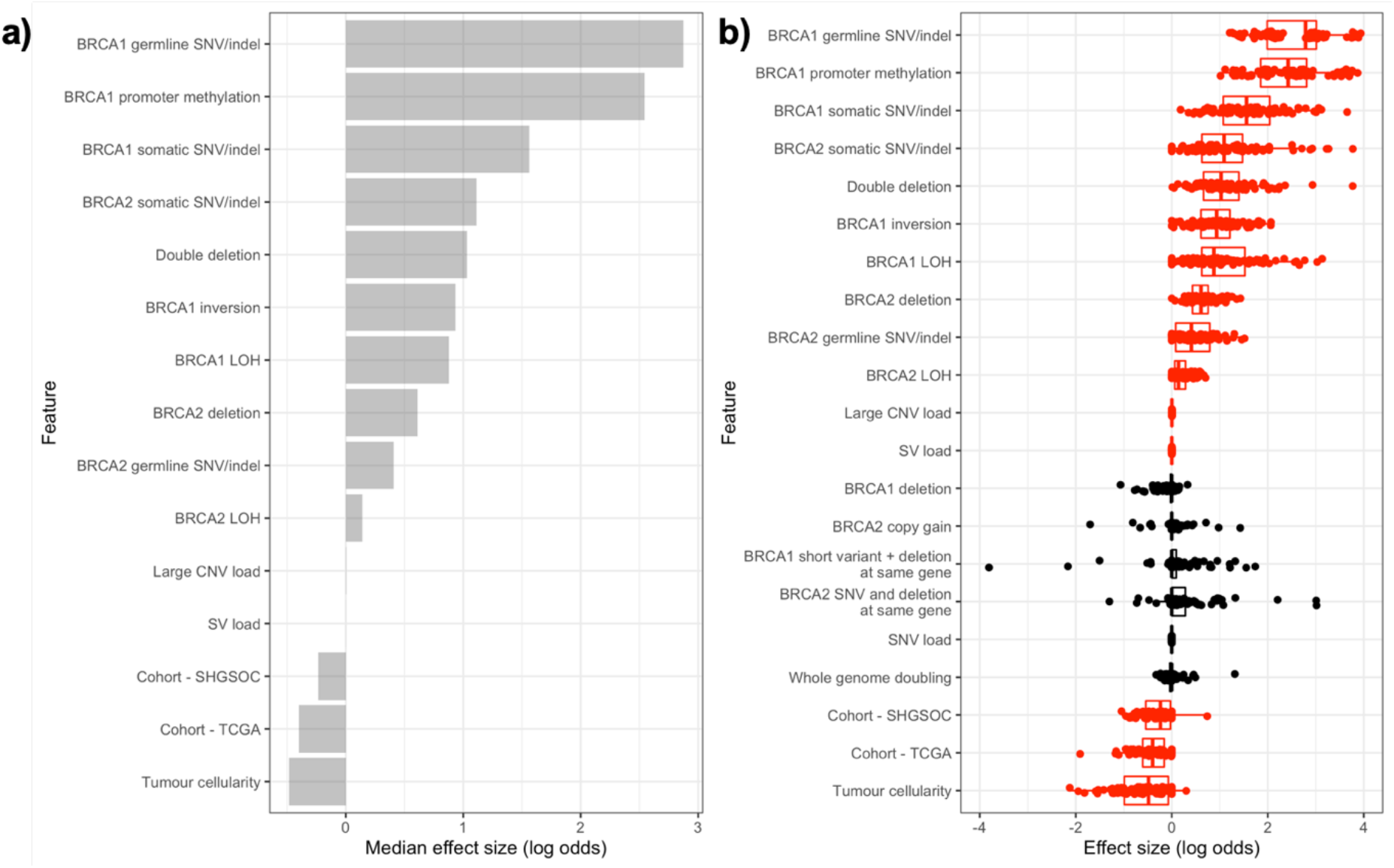
Integrative modelling of repair deficiency in HGSOC. a) Median effect sizes of genomic features selected to predict HRD, using elastic net regularised regression on 100 training/test set splits. Model performance was measured for each split and average AUC = 0.75. Binary mutational status variables (e.g. presence/absence of *BRCA1* somatic SNV) were included as factors and continuous variables were standardised to allow comparisons between variables. b) Distributions of effect size for each variable on HRD (log odds) in each training/test set split. Variables in red are selected for inclusion by the model in more than half of the training sets.

We can explain the observed pattern of HRD by the presence of mutational or epigenetic defects at the *BRCA1/2* genes in 81 out of 106 samples with predicted HRD (72 GSV/SSV/SV at *BRCA1/2*, 9 with *BRCA1* promoter methylation) but a further 25 samples with HRD remain unexplained. On further examination, we found that all of these samples harboured damaging GSVs and/or SSVs at other HR genes (defined by KEGG pathway annotation; Supplementary Tables 5, 6 and 7), motivating analysis of the potential roles of mutations at loci other than *BRCA1/2* and their inclusion in an expanded model. We also incorporated the combined expression levels of the genes dysregulated in the presence of a *BRCA1/2* short variant in the model as discussed in the previous section. However, due to the lower number of samples with expression information and the increased number of features this model is likely to be underpowered to accurately identify significant features and we found no convincing evidence for a strong influence of mutations at other HR genes or expression of genes other than *BRCA1/2* on HRD (Supplementary Figure 5). We conclude that the current gold-standard of genomic data for HGSOC supports the role of genomic events other than short variants at *BRCA1/2;* however, mutations at other HR genes are less informative for HRD prediction.

### HRD is associated with longer survival in the absence of disruptive short variants at *BRCA1/2*

HGSOC tumours with HRD show increased responsiveness to platinum agents, and a recent trial showed that the use of olaparib as first-line maintenance therapy in women with newly diagnosed advanced ovarian cancer and a *BRCA1/2* germline or somatic mutation led to a 70% lower risk of disease progression or death compared to placebo^36^. However, the relationship between HRD in HGSOC resulting from events other than disruption via short variants at *BRCA1/2*, and overall survival or response to treatment has been less clear. Studies have demonstrated that HRD as a result of amplification or disruption of genes other than *BRCA1/2* is associated with better prognosis or improved treatment response ^11,37–39^. Some studies have reported no survival advantage, or worse survival for patients with *BRCA1* promoter methylation^6,8,40^. However, more recent analysis of TCGA data^1^ supports an association between HRD and longer overall survival in HGSOC.

The probability of HRD is significantly associated with longer overall survival in our combined cohort (Hazard ratio=0.36, 95% CI 0.24–0.54, p-value = 8.4×10^-7^) and this effect is only slightly attenuated by adjustment for patient age and tumour stage at diagnosis (Figure 6a). Notably, this effect persists (HR=0.46, 95% CI 0.28-0.74, p-value=1.5×10^-3^) (Figure 6b) when we exclude the patients with *BRCA1/2* GSV/SSV, who are already known to have longer survival. We see a similar effect when we examine the effect of HRD on progression-free survival (PFS) (HR=0.55, 95% CI 0.36 – 0.85, p-value = 0.007) which also is robust to the exclusion of patients with *BRCA1/2* GSV/SSV (HR=0.58, 95% CI 0.36-0.95, p-value=0.03). Considering HR deficiency as a binary endpoint, as is more realistic in a clinical setting, we observe the same effect on overall survival but a much weaker association with progression-free survival (Figure 6c),d)), which appears stronger in the subset of the cohort without short variants at *BRCA1/2*, because the patients with HRD tumours in this subset had earlier stage disease than the patients with non-HRD tumours. These effects are consistent with large disruptive deletions at *BRCA1/2* affecting overall survival in addition to the survival benefit conferred by short variants at *BRCA1/2*. Overall, the effect of HRD on survival is consistent with suggestions that signatures of HRD are predictive biomarkers for platinum or PARP inhibitor response that could be used in addition to *BRCA1/2* mutations^41^. Furthermore, this translates into longer overall survival for the patient. Similar analyses in other cancer types where HRD has been detected may also be informative. This suggests that HRD whether arising from short variants, SVs or combinations of these mutations at *BRCA1/2*, represents a consistent tumour phenotype, targetable by therapies exploiting HRD. In the shorter term, patient stratification based on short variants could be improved by the addition of a broader range of mutational events.

**Figure 6:**
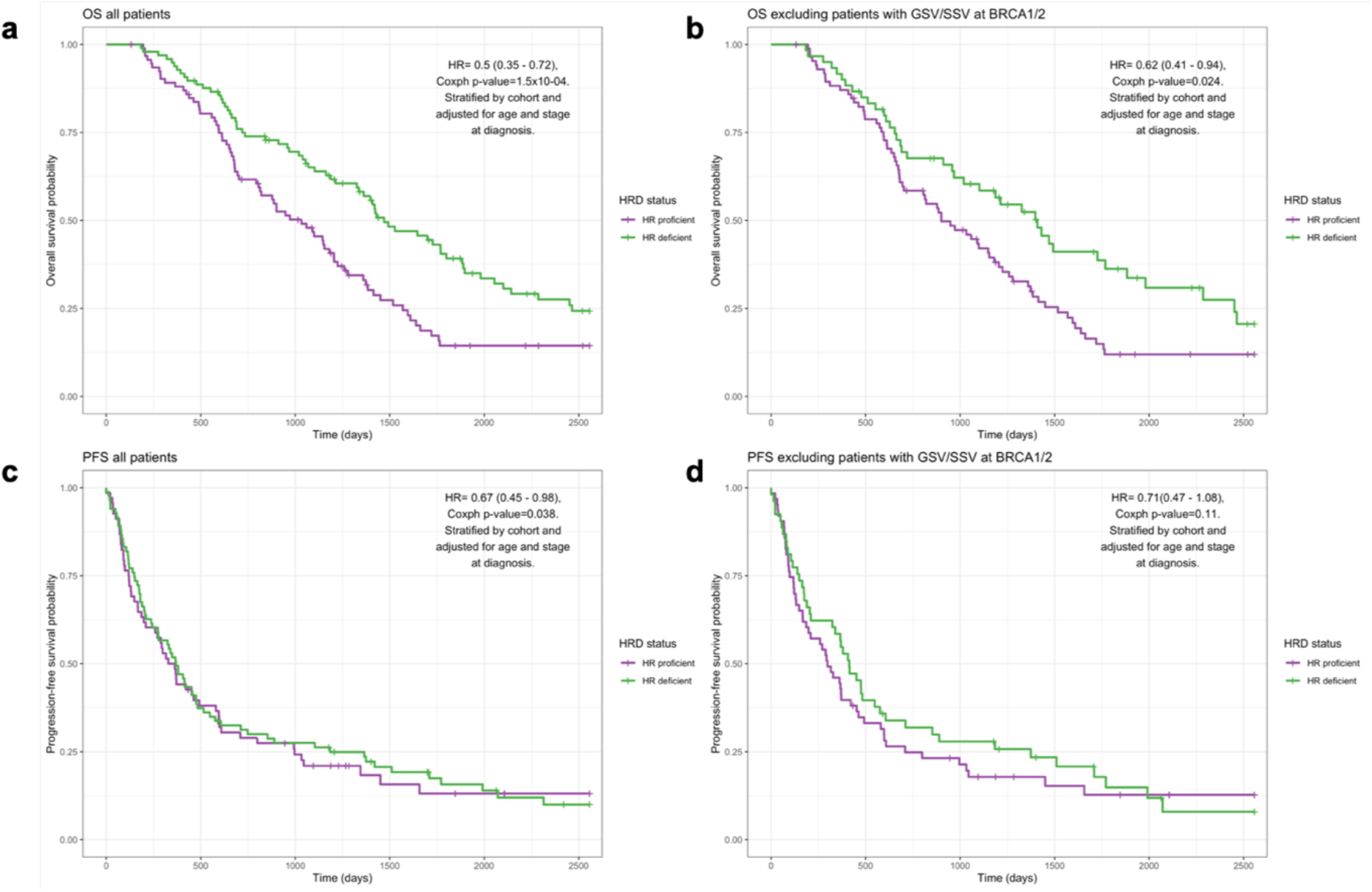
Predicted HRD is associated with patient survival in the absence of short variants at *BRCA1/2*. a) The effect of HRD on overall survival time after diagnosis (in days) in HGSOC. (N (events) =190 (144)). b) The effect of HRD on overall survival time after diagnosis (in days) in HGSOC patients without *BRCA1/2* GSV/SSV. (N (events) =145 (113)). c) The effect of HRD on progression-free survival-time after diagnosis (in days) in HGSOC. (N (events) = 151 (129)). d) The effect of HRD on progression-free survival-time after diagnosis (in days) in HGSOC. (N (events) = 115 (100)). Kaplan-Meier plots comparing survival between HR deficient and HR proficient patients as defined by HRDetect score above and below 0.7. Hazard ratio estimates taken from Cox proportional hazards models with HRDetect score as a covariate, stratified by cohort and adjusted for age and stage at diagnosis.

## Discussion

We have assembled the largest collection of HGSOC WGS data examined to date, with matched expression data for most of the 205 tumours included, revealing new insights into the genesis of HRD in HGSOC based upon genome-wide mutational spectra. We show that structural variation at *BRCA1/2* in HGSOC is frequent and is dominated by multi-megabase deletions encompassing the genes. Large deletions spanning *BRCA1/2* contribute to HRD independently of short variants, and samples with compound deletions affecting both *BRCA1* and *BRCA2* generate the highest risk of HRD. In addition, deletions and short variants contribute together to compound *BRCA1/2* genotypes with risk of HRD similar to the classical short variants known to be associated with HRD. Examining transcriptomic data for the same samples we have shown that large deletions overlapping *BRCA1/2* are associated with lower *BRCA1/2* expression, suggesting a direct impact of these deletions on gene function in many cases. The frequent inactivation of *BRCA1/2* by large deletions in HGSOC is novel to our knowledge, and the original analysis of the AOCS cohort reported only one *BRCA1/2* large deletion (>1Mb)^5^ (Patch et al (2015), Supplementary Table 4.2). The original TCGA cohort analysis did report frequent losses of the chr13q and chr17q chromosome arms (including the *BRCA1/2* loci) based upon SNP microarray data^6^ (Bell et al (2011), Supplementary Table S5.1), but such data are known to generate high levels of false positives and negatives^33,34^ and these losses were not postulated to affect *BRCA1/2* function. Thus previous assessments of SVs impacting the *BRCA1/2* loci have been characterised by under-reporting, likely to be a result of the use of less sensitive algorithms tuned to detect smaller focal deletions^5^ as well as CNA estimates derived from SNP microarray data^5,6^ and exome-restricted sequencing data^6,35^. Other types of structural variation are less frequent but still evident, such as large inversions at *BRCA1* and duplications at *BRCA2*. The impact of these categories of mutation on the function of the gene is less well studied but our data suggest that when *BRCA1* inversions in particular are considered together with the other mutational events in the tumour, their presence may aid prediction of HRD.

Finally, we have constructed an integrated model of HRD in HGSOC, including a large variety of mutation and expression-based variables across the combined cohort. This model supports an independent role for structural variation at *BRCA1/2* in HRD and highlights the diversity of routes that tumours may follow to reach HRD. Given this diversity, and the substantial fraction of samples where HRD is detected in the absence of any detectable *BRCA1/2* mutations, we conclude that the direct detection of HRD in HGSOC using genomewide sequencing data is a valuable addition to the search for inactivating mutations in HR pathway genes. This is likely to be the case for other cancers showing evidence for HRD, such as uterine, lung squamous, oesophageal, sarcoma, bladder, lung adenocarcinoma, head and neck, and gastric carcinomas^1^. The variety of events sufficient for a tumour to develop HRD is not well understood, but recent studies suggest that there is selective pressure for biallelic inactivation leading to HRD in cancer types with predisposing germline variants in the HR pathway, such as breast, ovarian, pancreatic and prostate cancers^36^.

One of the key challenges in studies of this type is deciding upon a ‘gold standard’ test of HRD. Current functional, clinical and molecular tests all have advantages and disadvantages. The limitations of HRDetect include that it was developed and trained using breast tumour data and is predicated on BRCA1/2 deficiency arising from SSV disruption and promoter methylation, rather than any form of disruption to any HR gene. Although in the context of the current study, ofBRCA1/2 disruption by SV, the latter is of less importance. All current genomic HRD tests are further limited to demonstrating that HRD once existed in the evolution of a tumour, and are blind to the restoration of HR by events such as secondary mutations, as well as hypomorphic HRD variants and epigenomic changes.

Accurately identifying tumours with HRD is crucial to predict PARP inhibitor sensitivity. PARP inhibitor maintenance studies in HGSOC have demonstrated an exceptional benefit in both the first line and relapsed disease settings^14,15,37–41^. In the *BRCA1/2* deficient subgroup, there is a 3-5 fold reduction in the risk of progression or death from the use of these agents. In patients with functional *BRCA1/2*, the HRD population is identified using commercially available assays such as the Myriad MyChoice assay (which creates a score based upon large-scale transitions, loss of heterozygosity and telomeric imbalance) or the Foundation Medicine LOH (loss of heterozygosity) assay. These assays provide some enrichment of patients who were PARP inhibitor sensitive but they are unable to identify patients who did not benefit from PARP inhibition^14,15,39^. HR proficient tumours have been defined as those tumours which harbour low genome-wide rates of mutation and lack inactivating *BRCA1/2* short variants and *BRCA1* promoter methylation^3^. There may be great impact in establishing which genomic features are present, rather than absent, in HR proficient tumours. In common with other studies, we compared the risk of HRD associated with various types of *BRCA1/2* mutation to samples lacking detectable *BRCA1/2* mutations instead of samples with known HR proficiency. This set is likely to include some hidden HRD samples and as a result, we expect our estimates of the effects of *BRCA1/2* disruption on the risk of HRD to be conservative.

There is an urgent clinical need to better understand the processes that give rise to both *BRCA1/2* loss and more broadly contribute to HRD. Our study demonstrates that *BRCA1/2* loss by structural variation may have a comparable impact on HRD and patient survival to short variants at *BRCA1/2*. However, these variants are unlikely to be detected by sequencing methods currently employed in the clinic.

## Data availability

Previously published WGS and RNA-seq data that were reanalysed here are available via EGA at accession code EGAS00001001692 (ICGC PCAWG). WGS, RNA-seq and clinical data from the Scottish cohort (SHGSOC) will be made available via EGA at accession code EGAS00001004410. Other supporting data have been provided in the Supplementary Tables.

## Code availability

All code will be made available at https://github.com/ailithewing.

## Acknowledgements and funding support

A.E is supported by a UKRI Innovation Fellowship (MR/RO26017/1). C.A.S, A.Me and G.R.G are supported by MRC core funding to the MRC Human Genetics Unit (MRC grant MC_UU_00007/16). R.L.H is supported by an MRC-funded Research Fellowship. S.D received funding from AstraZeneca and the Beatson Cancer Charity. I.A.McN acknowledges funding from Ovarian Cancer Action and the NIHR Imperial Biomedical Research Centre. A.V.B acknowledges funding from Cancer Research UK (C29717/A17263, C29717/A18484, C596/A18076, C596/A20921, A23526), Wellcome Trust Senior Investigator Award (103721/Z/14/Z), Pancreatic Cancer UK Future Research Leaders Fund (FLF2015_04_Glasgow), MRC/EPSRC Glasgow Molecular Pathology Node, The Howat Foundation.

Sequencing of the SHGSOC cohort was supported by AstraZeneca, the Medical Research Council and the Scottish Chief Scientist through a Precision Medicine Scotland Innovation Centre/Scottish Genome Partnership (SEHHD-CSO 1175759/2158447) collaboration. This Scottish Genomes Partnership is funded by the Chief Scientist Office of the Scottish Government Health Directorates [SGP/1] and The Medical Research Council Whole Genome Sequencing for Health and Wealth Initiative (MC/PC/15080). This study would not be possible without the families, patients, clinicians, nurses, research scientists, laboratory staff, informaticians and the wider Scottish Genomes Partnership team to whom we give grateful thanks. The authors would also like to acknowledge the Edinburgh Clinical Research Facility for the sequencing of RNA samples from the SHGSOC cohort.

The authors would also like to extend our thanks to the Nicola Murray Foundation, and the Edinburgh Ovarian Cancer Database, from which the clinical data for much of the Scottish cohort were retrieved. We also thank the NRS Lothian Human Annotated Bioresource, NHS Lothian Department of Pathology, the Edinburgh Experimental Cancer Medicine Centre, the Biorepository at the Glasgow Queen Elizabeth University Hospital and the Tayside Biorepository for their support.

## Author contributions

C.G, C.A.S, C.S.H, I.A.M, T.M, M.F, N.M, A.M, B.D, R.M, J.C.B, A.V.B conceived the study within the broader remit of the Scottish molecular ovarian cancer collaboration. A.E, C.A.S, C.G conceived and designed the analysis of this cohort. M.C, R.L.H, M.F, M.L, T.M, N.M, N.S,

S.D, M.M, F.N, R.G, P.R and C.G acquired patient samples. C.S.H performed histopathological review. M.C, I.C and B.D performed sample processing. B.M and A.V.B performed whole genome sequencing. A.Me, A.E, G.R.G processed the sequencing data. T.R, R.L.H, C.B, T.M, M.F, N.M, M.L, S.D. R.G, M.M, F.N, P.R and C.G provided clinical data. A.E performed statistical analyses. P.R, C.G, C.A.S, L.M, A.M, B.D, R.M, J.C.B and A.V.B provided strategic direction to the project. SGP and AstraZeneca funded the work. A.E, C.A.S and C.G drafted the manuscript. All authors read and commented on the manuscript and approved the final version.

## Methods

### Scottish sample collection and preparation for WGS

Scottish HGSOC samples were collected via local Bioresource facilities at Edinburgh, Glasgow, Dundee and Aberdeen and stored in liquid Nitrogen until required. HGSOC patients were determined from pathology records and were included in the study where there was matched tumour and whole blood samples. On receipt of tumour material the tumour was processed as follows: firstly, the tumour sample was divided into two for DNA and RNA extraction. Slivers of tissue were cut from the front and rear faces of the DNA sample, then fixed in formalin and embedded in paraffin wax. Sections from the front and rear tissues from all samples were examined by H&E staining supplemented by WT1/p53 immunohistochemistry if required. Following pathology review, samples were only included if they met the following criteria: they were confirmed as HGSOC and there was greater than 40% tumour cellularity throughout the tumour, determined using the H&E sections. Somatic DNA was extracted using the Qiagen DNeasy Blood and tissue kit (cat no 69504). The tissue was initially homogenised using a Qiagen Bioruptor, followed by the manufacturers recommended protocol (including RNase digestion step). Germline DNA was extracted from 1-3ml whole blood using the Qiagen FlexiGene kit (cat no 51206) following the manufacturers recommended protocol. The resulting DNA underwent quality control as follows: firstly, A260 and A280nm were measured on a Denovix DS-11 Fx to qualitatively illustrate A260/280nm and A260/230nm ratios as surrogate measures of DNA purity. A260/280 had to be 1.8 or greater and A260/230 had to be 2.0 or greater. Then, DNA was quantified using LifeTechnologies Qubit dsDNA BR kit (cat no Q32850) and we required a minimum of 50ul at 25ng/ul for WGS. Thirdly, DNA was diluted to 25ng/ul and a representative sample was loaded onto a 0.8% TAE gel, ran at 100v for 60mins and then imaged using a BioRad ChemiDoc imaging system to visualise the DNA quality. Only when all 4 quality control requirements were satisfied was the DNA sequenced. The DNA was sequenced at the Glasgow Precision Oncology Laboratories.

### Sequence acquisition

WGS and RNA-seq reads were downloaded in compressed FASTQ format from the sequencing facility (SHGSOC) or in aligned BAM format (including unaligned reads) from the European Genome/Phenome Archive (AOCS) and the Bionimbus Protected Data Cloud (TCGA_US_OV). The reads obtained in BAM format were query-sorted using sambamba 0.6.8^42^ and converted to FASTQ with bamtofastq from the biobambam2 2.0.87 package^43^.

### Primary processing of WGS

Reads were aligned to the hg38 reference genome using a bcbio 1.0.7 pipeline^44^ with bwa 0.7.17 aligner^45^ and Genome Analysis Toolkit 4.0.0.0^46,47^ post-processing (see Supplementary Methods for full pipeline configuration, program and resource versions).

Somatic and germline variant calling was also run with a bcbio 1.0.7 pipeline^44^. Germline SNPs and indels were called with GATK 4.0.0.0 HaplotypeCaller^46,47^. Somatic SNVs and indels were called as a majority vote between Mutect2^48^, Strelka2^49^ and VarDict^50^. Small variants were annotated with Ensembl Variant Effect Predictor v91^51^ and filtered for oxidation artifacts by GATK 4.0.0.0 FilterByOrientationBias^46,47^. Somatic structural variants were called with Manta 1.2.1^52^ and somatic copy number variants with CNVkit 0.9.2a0^53^. Loss of heterozygosity and somatic copy number variants were also identified with CLImAT^54^. Whole genome duplication estimates were based on allele-specific somatic CNVs called by FACETS^55^. Structural and copy number variants were annotated with Ensembl Variant Effect Predictor v91^51^. Sample quality control was performed with Qsignature 0.1^56^ to identify sample mix-ups and VerifyBamId 1.1.3^57^ to identify sample contamination. Tumour cellularity was estimated using both CLImAT’s estimates and p53 variant allele frequency. These measures were compared to the qPure estimates for the AOCS cohort^5^ and histopathological estimates for the SHGSOC cohort with very good concordance (Supplementary Figure 6). The CLImAT estimates were used as the final estimates of cellularity.

### Filtering of small variants (SNVs and indels) at *BRCA1/2*

Germline short variants at *BRCA1/2* were filtered to include only damaging pathogenic variants for the purposes of establishing *BRCA1/2* mutational status. Included variants were all of moderate or high impact according to VEP^51^. Variants with a pathogenic or risk factor annotation according to ClinVar^58^ were included (n=145). Remaining variants with a ClinVar benign or likely benign status were excluded (n=1147). Remaining frameshift or nonsense (stop gained) or splice donor/acceptor variants were included (n=125). Remaining missense variants with damaging SIFT^59^ and PolyPhen^60^ predictions were included (n=36). Remaining missense variants called as damaging by only one of SIFT and PolyPhen were considered borderline and were excluded if their CADD score < 20^61^. Missense or inframe variants with no Clinvar, SIFT or PolyPhen evidence were excluded.

Somatic short variants at *BRCA1/2* were also filtered for pathogenicity to include variants that: were annotated by VEP as being of high or moderate impact, were pathogenic according to at least one of SIFT or PolyPhen and had a high CADD score. In addition, we excluded somatic variants with an allele frequency less than 0.4.

### Curating a high-confidence list of structural variants at *BRCA1/2*

We identified structural variants in HGSOC patients using Illumina’s paired and split read based structural variant detection tool, Manta^52^. The ICGC-TCGA DREAM Somatic Mutation Calling Challenge (SMC-DNA)^22^ found that Manta performed better when scored using the harmonic mean of its precision and recall than taking an ensemble approach of structural variant callers. However, we observed that Manta was failing to detect a large number of very large deletions (>1Mb) that had been identified using depth of coverage-based approaches in PCAWG^62^. It is unsurprising that CNVs of this size are being missed as identifying them using split and paired read technologies will be hard in some cases and impossible in others. Therefore, we chose to supplement these calls with copy number variants greater than 1 Mb in size that were called by one caller (CNVkit) using evidence from read depth and were also confirmed by an allele-specific copy number caller (CLImAT). CLImaT provides an additional layer of evidence in addition to read depth as it also incorporates the shift in allele frequency of heterozygous SNPs within the potential copy number variant into its variant calling algorithm which also accounts for aneuploidy and sample cellularity. Deletions were assumed to be heterozygous if the copy number as estimated by both CNVkit and CLImAT was 1. It is possible that these deletions are homozygous deletions in a subclone of the tumour but as we expect that these events are early events^63^ we believe that it is more likely that they are true heterozygotes so have assumed this more conservative estimate of allelic loss. We visually inspected all the identified structural variants in the Integrative Genomics Viewer (IGV) (v2.4.10)^64^ using a log2 coverage bigWig file generated using bamCompare (v3.20) from the deeptools suite^65^. The bigWig file compared coverage between tumour and normal pairs normalized for sequencing depth. The magnitude of coverage and log fold change was inspected to confirm either duplication or deletion. For large SVs (>30kb) a chromosome wide view of the log2 track was considered. For structural variants in the range 300bp-30kb, the paired end sequencing reads were manually reviewed including looking at split reads, paired end insert size, read coverage and pair-orientation. We compared our filtered set of large CNVs in the samples included in PCAWG to PCAWG’s copy number calls and found that 40/41 of our variants in these samples were also identified by PCAWG.

### Implementation of HRDetect

To predict the level of HR deficiency in each tumour sample we implemented the HRDetect algorithm as published by Davies and Glodzik et al^3^. The algorithm is a logistic regression model with the probability of HR deficiency defined as ‘BRCA-ness’ as the outcome. The variables that make up the linear predictor represent genomic signatures that have been shown to correlate well with *BRCA1/2* mutation status. They include: the proportion of indels with microhomology at the breakpoints; the contribution of COSMIC SNV signatures 3 and 8 to the mutational profile of the tumour; the contribution of rearrangement signatures 3 and 5 to structural variation in the tumour; and the value of an earlier predictor of HR deficiency, the HRDIndex^66^, which combines levels of genome-wide medium length runs of loss of heterozygosity (LOH), telomere allelic imbalance (TAI) and large state transitions (LST). We based our implementation on a Snakemake pipeline made publicly available by Zhao et al^67^, with some modifications to ensure accurate recapitulation of the original method. As some of the AOCS cohort included here were also used in the validation of HRDetect in the original publication we were able to compare our implementation for the same patients with that of the authors.

Zhao’s pipeline makes use of the R package HRDtools^67^ in order to determine the value of the HRIndex. We used the same method determined by Zhao et al with the exception that we took the mean of the three inputs (LOH, TAI and LST) instead of the sum to reflect the original HRDetect approach. In addition, we redefined microhomology at indel breakpoints as an overlap between the deletion and the flanking region that is less than the full length of the deletion. In order to determine the contribution of each of the signatures to the mutational profile of each tumour we implemented three different methods: deconstructSigs^68^, SignIT^69^ and SigProfilerSingleSample^70^. Ultimately, we chose to use deconstructSigs as its estimates were the most strongly correlated with the results from the original HRDetect paper.

We used the weights for the independent variables that were defined by the original model rather than retraining the model on our data as the original weights trained on a breast cancer dataset have been shown to perform well on ovarian cancer datasets and our total sample size is substantially smaller than that used to train the model originally. As input we used the somatic SNV and indel calls identified by the ensemble calling approach described above; the structural variant calls made by Manta and the copy number segments defined by CNVkit. After determining the value of each of the components of the linear predictor for each sample, each of these input variables were standardized using the corresponding mean and standard deviation for the variable in question in the dataset that was used to determine the weights in the original model. Our implementation of HRDetect was very highly correlated with the original HRDetect implementation on the same samples (Spearman’s rho = 0.92).

### Scottish RNA sample preparation and sequencing

HGSOC samples were collected and underwent quality control as described for the DNA samples used for WGS. Somatic RNA was extracted from the resulting RNA sample using the Qiagen Qiasymphony RNA protcol (cat no 931636). The tissue was initially homogenised using a Qiagen Bioruptor, followed by the manufacturers recommended protocol (including DNase digestion). The resulting RNA the underwent quality control as follows: firstly, A260 and A280nm were measured on a Denovix DS-11 Fx to qualitatively illustrate A260/280nm and A260/230nm ratios as measures of RNA purity. A260/280 had to be 2.0 and A260/230 had to be 2.0-2.2. Then RNA was quantified using LifeTechnologies Qubit RNA BR kit (cat no Q10210). RNAseq was carried out by the Edinburgh Clinical Research Facility on an Illumina NExtSeq500 as detailed below.

Total RNA samples were assessed on the Agilent Bioanalyser (Agilent Technologies, #G2939AA) with the RNA 6000 Nano Kit (#5067-1512) for quality and integrity of total RNA, and then quantified using the Qubit 2.0 Fluorometer (Thermo Fisher Scientific Inc, #Q32866) and the Qubit RNA HS assay kit (#Q32855). Libraries were prepared from total-RNA sample using the NEBNext Ultra 2 Directional RNA library prep kit for Illumina (#E7760S) with the NEBNext rRNA Depletion kit (#E6310) according to the provided protocol. 400ng of total-RNA was then added to the ribosomal RNA (rRNA) depletion reaction using the NEBNext rRNA depletion kit (Human/mouse/rat) (#E6310). This step uses specific probes that bind to the rRNA in order to cleave it. rRNA-depleted RNA was then DNase treated and purified using Agencourt RNAClean XP beads (Beckman Coulter Inc, #66514). RNA was then fragmented using random primers before undergoing first strand and second strand synthesis to create cDNA. cDNA was end repaired before ligation of sequencing adapters, and libraries were enriched by PCR using the NEBNext Multiplex oligos for Illumina set 1 and 2 (#E7500). Final libraries had an average peak size of 271bp. Libraries were quantified by fluorometry using the Qubit dsDNA HS assay and assessed for quality and fragment size using the Agilent Bioanalyser with the DNA HS Kit (#5067-4626). Sequencing was performed using the NextSeq 500/550 High-Output v2 (150 cycle) Kit (# FC-404-2002) on the NextSeq 550 platform (Illumina Inc, #SY-415-1002). Libraries were combined in an equimolar pool based on the library quantification results and run across 5 High-Output Flow Cell v2.5.

### Primary processing of RNA-seq

RNA-seq data was analysed with bcbio-nextgen v1.0.8^44^ using the Illumina RNA-seq (https://github.com/bcbio/bcbio-nextgen/blob/master/config/templates/illumina-rnaseq.yaml) best practice template. Briefly, reads were aligned to the hg38 reference genome using hisat2^71^. Quality control was carried out using FastQC v0.11.8, FeatureCounts v1.6.4^72^, Qualimap v2.2.2-dev^73^ and reporting done using MultiQC v1.7^74^. Salmon quant v0.12.0^75^ was used to quantify the expression of transcripts against the hg38 RefSeq transcript database indexed using the salmon index (k-mers of length 31). For the AOCS and TCGA cohorts (paired-unstranded) salmon quant was run with the following options (-IU) and for SHGSOC (paired-stranded) the following options were used (-ISR). The tximport package v1.12.1^76^ was loaded from Bioconductor (Release 3.9)^77,78^ for use in R (v3.6.0) to import and summarize salmon transcript-level abundance estimates for further gene-level analyses. For differential expression analyses, expression counts were loaded directly from tximport into the DESeq2 package v1.24.0^79^. For visualization of gene expression, counts were normalized using the variance stabilizing transformation.

To further explore the functional impact of *BRCA1/2* mutations we collated previously published RNA-seq data available for the AOCS^5^ (N=80) and TCGA^6^ (N=30) cohorts, together with novel RNA-seq data for the SHGSOC (N=40) cohort generated for the present study as detailed above.

### Curation and acquisition of the patient’s clinical information

#### Scottish High Grade Serous Ovarian Cancer (SHGSOC)

Clinical data for the SHGSOC cohort was retrieved from the Edinburgh Ovarian Cancer Database^80^, the CRUK Clinical Trials Unit Glasgow and available electronic health records (ethics reference 15/ES/0094-SR751).

#### Australian Ovarian Cancer Study (AOCS) & The Cancer Genome Atlas (TCGA)

The clinical information including survival end-points, age and stage at diagnosis is available for these patients as part of the PCAWG project^21^.

### Statistical analyses

All downstream statistical analyses were carried out in R (v3.6.0) using Jupyter notebook (v4.3.1).

### Enrichment of large deletions at BRCA1/2 in HRD samples

Circularised permutation was carried out using the R package RegioneR^81^ to investigate whether large deletions overlap more often with *BRCA1* and *BRCA2* than they do elsewhere in the genome. We carried out 1,000,000 permutations to simulate the null hypothesis for each gene and judged significance at alpha = 0.05. (Supplementary Figure 2).

### Univariable analyses of genomic features and risk of HR deficiency

The risk of HR deficiency in tumours with *BRCA1/2* mutations, grouped by type, relative to those tumours without *BRCA1/2* mutations were determined using Fisher’s exact tests. The effect of mutations at *BRCA1* and *BRCA2* were determined together and, where sample size permitted, separately for distinct mutually exclusive mutation categories including: germline short variants only, somatic short variants only, single deletion in the absence of a short variant or other SVs, deletion of both *BRCA1* and *BRCA2* in the absence of a short variant, single non-deleting SV in the absence of short variants or deletion. We also considered the impact of the presence of a short variant accompanied by a deletion at either of the genes. The relative risk conferred by each mutational category was calculated in comparison to the group of patients without *BRCA1/2* mutations. Samples where BRCA1/2 promoter methylation had been detected were excluded except for where the effect of BRCA1 promoter methylation was itself being examined. All samples with BRCA1 promoter methylation are HR deficient so pseudo counts of 1 are used to estimate the effect size which is therefore a likely underestimate. P-values were adjusted for the impact of multiple testing using Benjamini-Hochberg correction and were considered together with the effect sizes in the reporting of results.

### Differential expression analyses of *BRCA1/2* in tumours with and without deletions at *BRCA1/2*

In order to compare the gene expression levels at *BRCA1/2* between tumours with and without *BRCA1/2* deletions, we used the package DESeq2 to test for differential expression between the raw gene expression counts at each gene between samples with and without a deletion at that gene. At *BRCA1*, samples that also had a short variant had significantly lower expression than those that had a deletion alone. This was driven by SNVs resulting in a stop gain variant and the presence of indels. As a result we only considered the samples with a deletion in the absence of a short variant. At *BRCA2*, this was not the case and the samples with short variants in addition to a deletion had comparable levels of expression to those with deletions alone so were included in the analysis. Cohort and tumour sample cellularity were included as covariates in the model formula.

### Identifying differentially expressed genes in the presence of HRD

For the samples with RNAseq information, we defined a conservative HR deficient group which included the samples with pathogenic short variants at *BRCA1/2* either in the germline or in the tumour (N=50). The contrasting HR proficient group of tumours, consisted of samples without damaging *BRCA1/2* short variants or *BRCA1* promoter hypermethylation or damaging short variants at HR genes as defined by KEGG and a quiet mutational profile defined by absence of the HRD related rearrangement signatures (N=47). This is consistent with the definition of HR proficiency used to train HRDetect. We used DESeq2 to compare the expression of all protein coding genes between the two groups and identified those genes that were differentially expressed. Cohort and tumour cellularity were included as covariates in the model. We used a log fold change threshold of 1 and a Benjamini-Hochberg adjusted p-value threshold of 0.05 to indicate significant differential expression. Functional annotation of the differentially expressed genes was carried out by comparing the differentially expressed genes with a background list of all protein-coding genes and testing for enrichment of the differentially expressed genes in curated gene lists from GO: BP, CC and MF and KEGG. This was done using clusterProfiler^82^with p-value and q-value thresholds of 0.05.

We defined a gene expression signature for HR deficiency by running principal component analysis on the variance stabilising transformed counts of the differentially expressed genes using all of the samples. The first principal component was taken as the gene expression signature for HRD with HR deficient samples having significantly lower values of the signature than HR proficient samples. We tested whether HR deficient and proficient samples had significantly different levels of the signature using a Wilcoxon Rank Sum test (Mann-Whitney U test).

The ability of a gene expression signature for HRD to predict HRD was assessed by identifying differentially expressed genes between only 80% of true HR deficient and 80% of true HR proficient samples and examining whether the HR deficient and proficient samples in the test set lay at significantly different points along the main axis of variation (first principal component) in the expression of these genes within the test set. The difference in the levels of the signature for HR deficient and proficient samples within the test set was tested using a Wilcoxon Rank Sum test.

### Multivariable elastic-net regularised regression model

Given the relative sparsity of the data and the correlation between features we used a multivariable elastic-net regularised regression model for the binary outcome of HRD defined by a probability of HRD greater than 0.7 from HRDetect. The model was implemented using the glmnet^83,84^ package in R. The data were partitioned into train and test sets (80:20) and the tuning parameters were optimised, in order to maximise the AUC, using 10-fold cross validation of the training set. The model was then fitted to the training set and the model performance assessed using the test set. The input variables available for selection were: *BRCA1* germline short variant status; *BRCA1* somatic short variant status; the presence of a large somatic deletion at *BRCA1;* the presence of a large somatic deletion at *BRCA1* and a *BRCA1* short variant; all the corresponding variables for *BRCA2;* the presence of an inversion at *BRCA1;* the presence of a duplication at *BRCA2;* the presence of a large somatic deletion at *BRCA1* and at *BRCA2* (double deletion); *BRCA1* promoter hypermethylation; whole genome doubling; genome-wide load of SNVs, large CNVs and SVs in addition to cohort and tumour cellularity. The data was partitioned and the model optimised and fitted to 100 train-test splits of the data in order to assess the robustness of the feature selection.

The model was then extended to include: pathogenic short variants in the germline or tumour at the HR genes defined by KEGG; the presence and load of large CNVs at the HR genes defined by KEGG; expression of *BRCA1* and *BRCA2;* the HRD gene signature; genomewide SNV load, SV load and large CNV load; sample cellularity and cohort. The gene expression signature for HRD was defined using the method described above but with differentially expressed genes determined specifically in each variation of the training set in order to avoid over-fitting.

### Survival-time analyses of the impact of HRD on overall survival

Follow up information including overall survival time was available for 190 out of 205 patients, of which 144 were deceased by the time of last follow up. The association between genome-wide patterns of HRD and progression-free survival was also assessed. Progression-free interval time was available for 151 of the patients from the AOCS and SHGSOC cohorts of which 129 relapsed by the time of last follow up. The effect of the HRDetect score, as a measure of the probability of HRD in the tumour, on the length of time that patients survived after diagnosis (overall survival-time) and the time between diagnosis and first radiologically defined progression (progression-free survival-time) was assessed using Cox proportional hazards models stratified by cohort. Multivariable models were also fitted adjusting for age and stage at diagnosis and the Schoenfeld residuals were examined. Survival probability through time was compared between HR deficient (HRDetect score > 0.7) and HR proficient (HRDetect score <=0.7) patients in Kaplan-Meier plots. This was repeated excluding the patients with *BRCA1/2* short variants to assess the impact of HRD driven by other events on survival.

## Structural variants at the BRCA1/2 loci are a common source of homologous repair deficiency in high grade serous ovarian carcinoma

### Supplementary figures

**Supplementary Figure 1:**
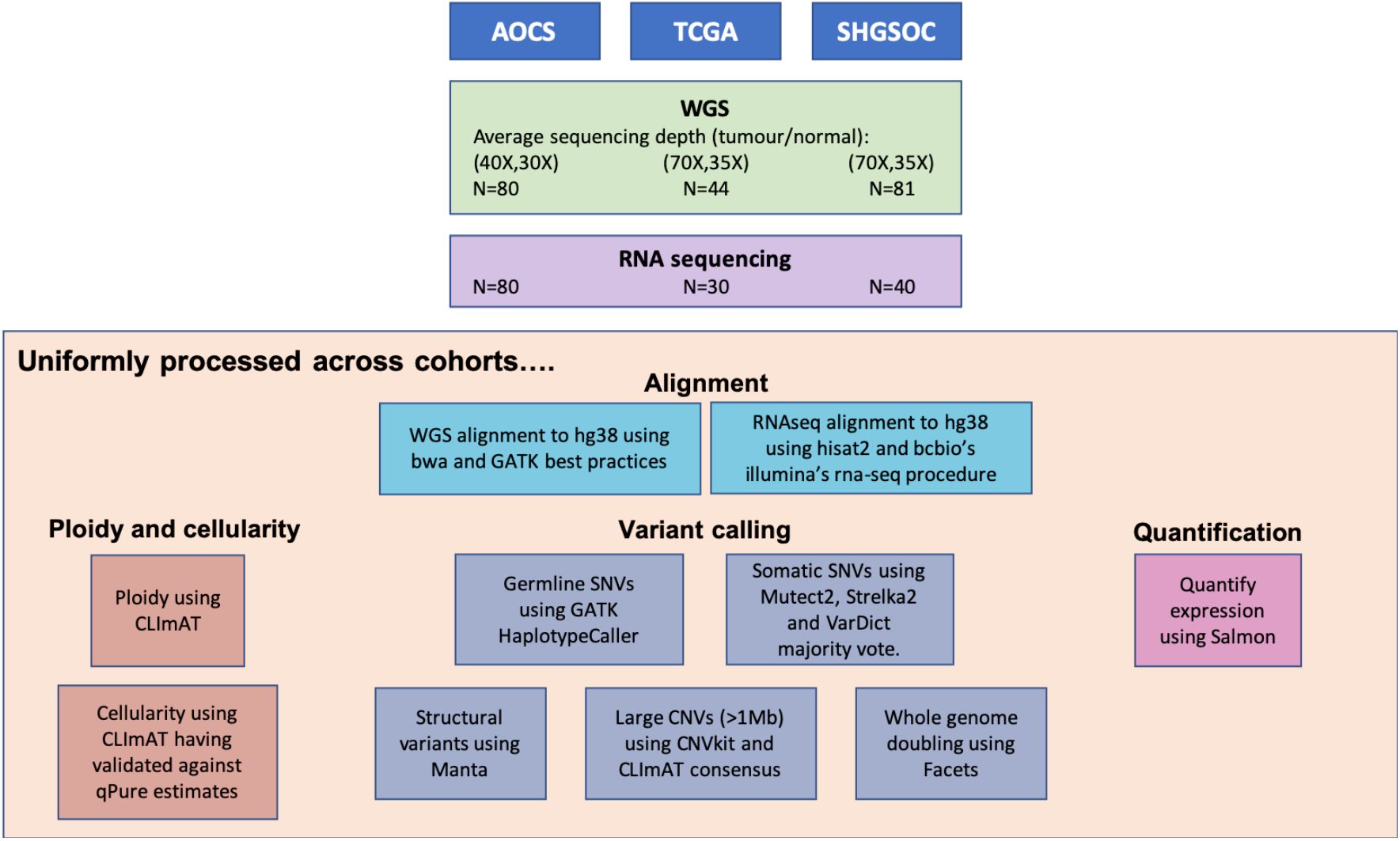
Uniform primary processing of three large HGSOC cohorts. WGS and RNA-seq fastqs were downloaded for AOCS and TCGA and the SHGSOC cohort was sequenced for the first time. Sequencing reads were aligned uniformly for all cohorts to hg38 and variant detection was carried out to detect a range of types of variant using existing published tools. Ploidy and cellularity were estimated using the allele-specific copy number caller CLImAT and gene-level expression counts were quantified using Salmon.

**Supplementary Figure 2.**
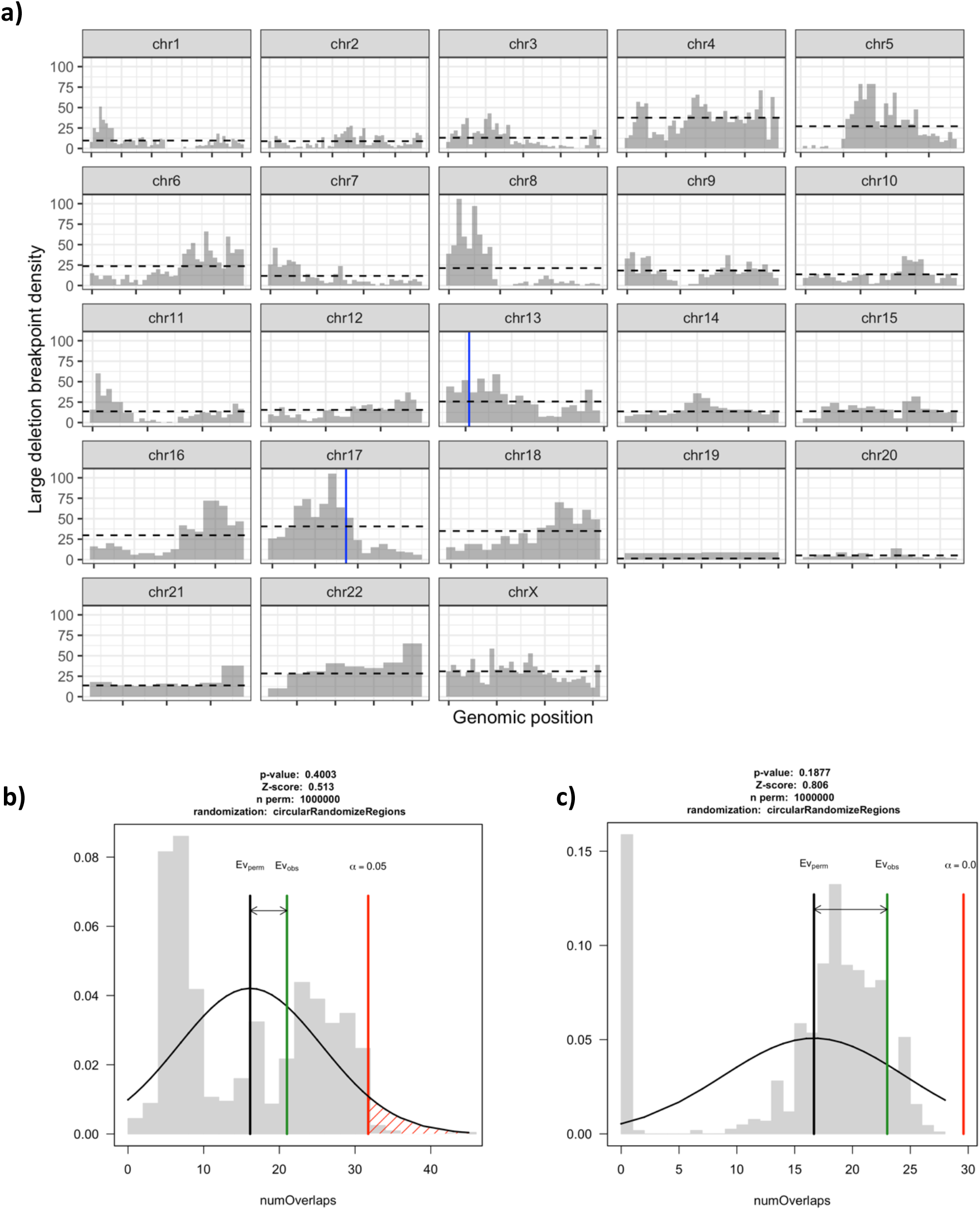
Enrichment testing of large deletions at BRCA1/2 in HRD samples. a) Pile up plot of large deletion breakpoints in HRD samples by genomic position. BRCA1 and BRCA2 are indicated by blue bars. b) Results of enrichment testing using circularised permutation to test whether *BRCA1* is enriched for large deletions in HRD samples relative to the rest of the genome. The green line indicates the number of large deletions overlapping *BRCA1* in HRD samples which is not significantly higher than elsewhere in the genome. c) The same but for *BRCA2* which also is not significantly enriched for large deletions in HRD samples.

**Supplementary Figure 3.**
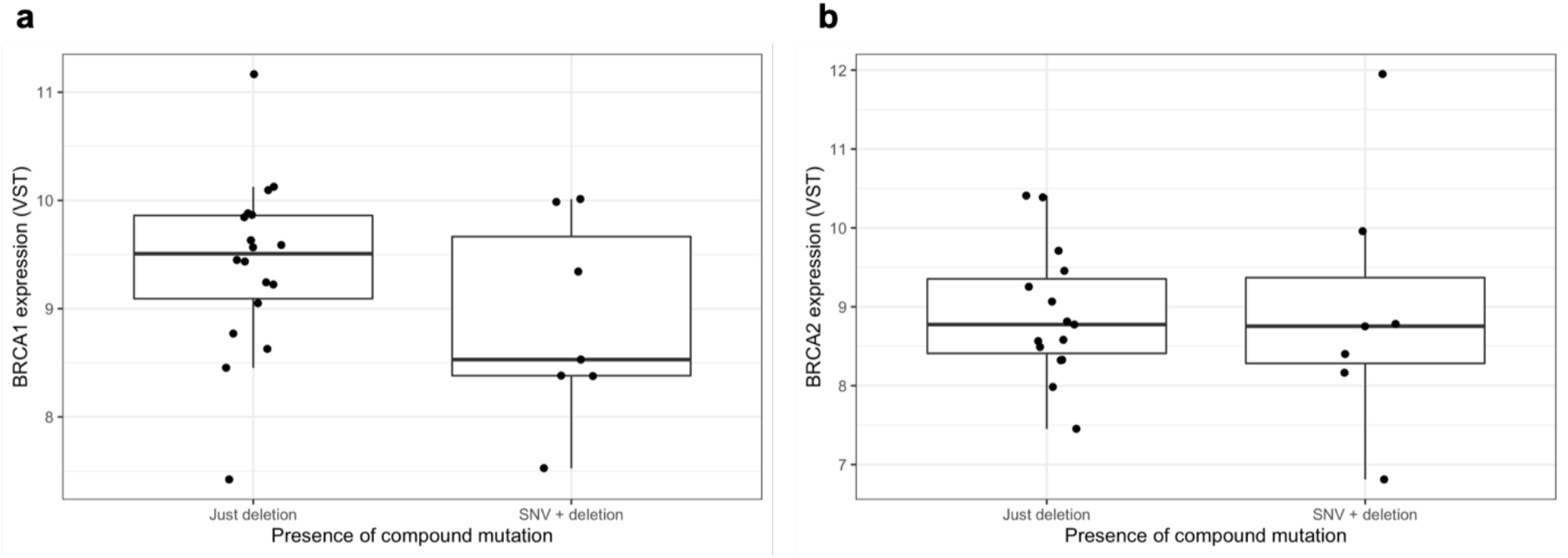
BRCA1/2 expression in samples with deletions, with and without SNVs in the same BRCA1/2 gene. a) Boxplot with points overlayed showing that BRCA1 expression (variance stabilising transformed) is higher in samples with only deletions than in samples with an SNV and a deletion at BRCA1 (DESeq2 fold change for just deletions vs SNV + deletion = 1.6, p-value=0.02). b) Boxplot with points overlayed showing no evidence of a significant difference in BRCA2 expression between samples with only deletions and samples with an SNV and a deletion at the same gene (DESeq2 fold change for just deletions vs SNV + deletion= 1.02, p-value=0.95).

**Supplementary Figure 4.**
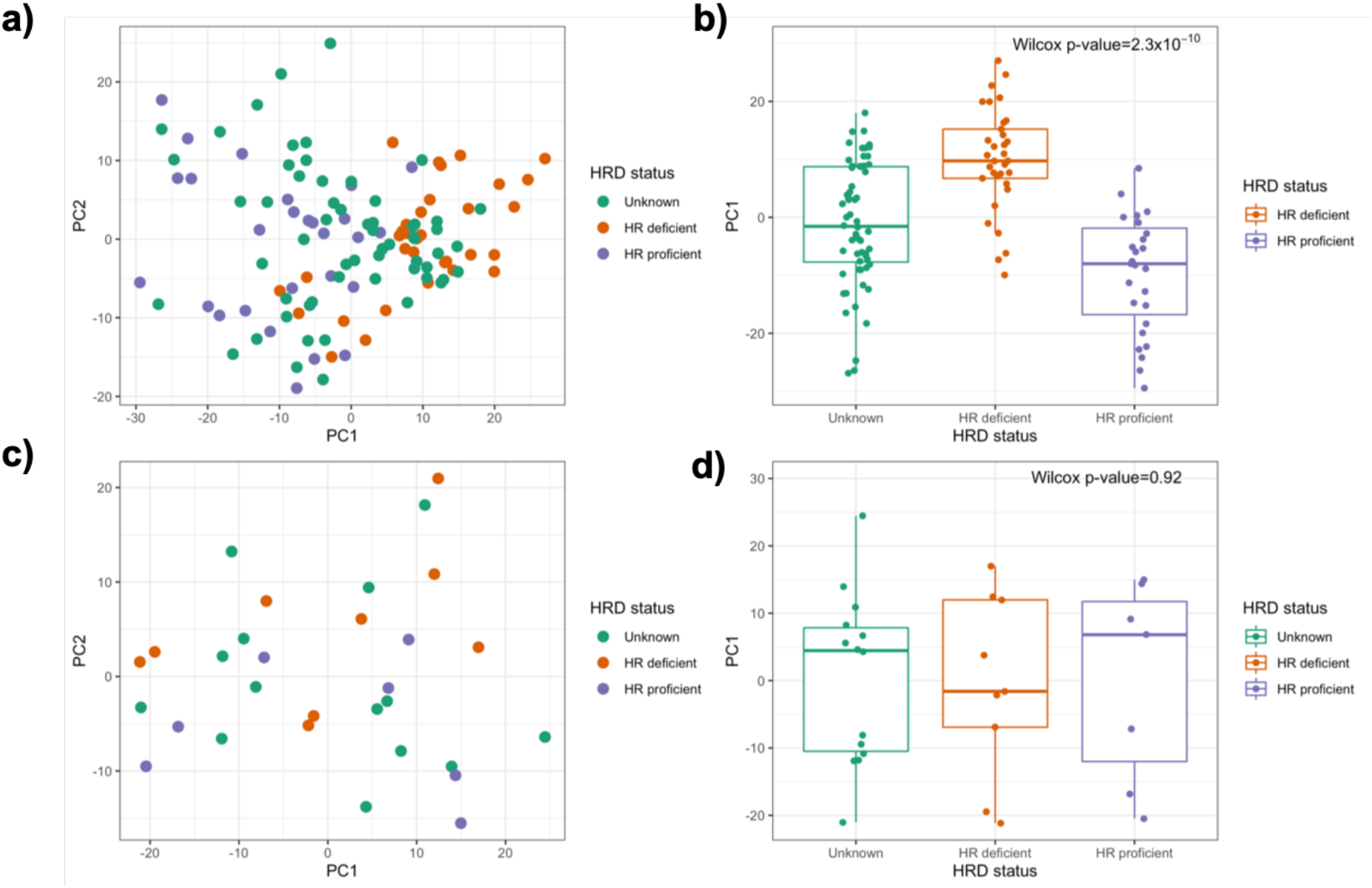
Performance of expression signature for HRD at predicting HRD. a) The first two principal components of differentially expressed (DE) genes between HRD and HRP samples. DE genes identified in the training set and PCA fitted to the training set. b) The level of the first principal component in samples in the training set. The first principal component, is significantly different between HR deficient and HR proficient samples in the combined cohort (Wilcox p-values =2.3×10^-10^) c) The first two principal components from PCA applied to the test set using the DE genes identified in the training set. d) The level of the first principal component in samples in the test set. PC1 discriminates poorly between HRD and HRP samples which suggests that HRD expression signatures is not a generalisable predictor. Notably these genes do not include known HR genes and given their diverse functions their dysregulation is likely to be a consequence rather than a cause of HRD.

**Supplementary Figure 5.**
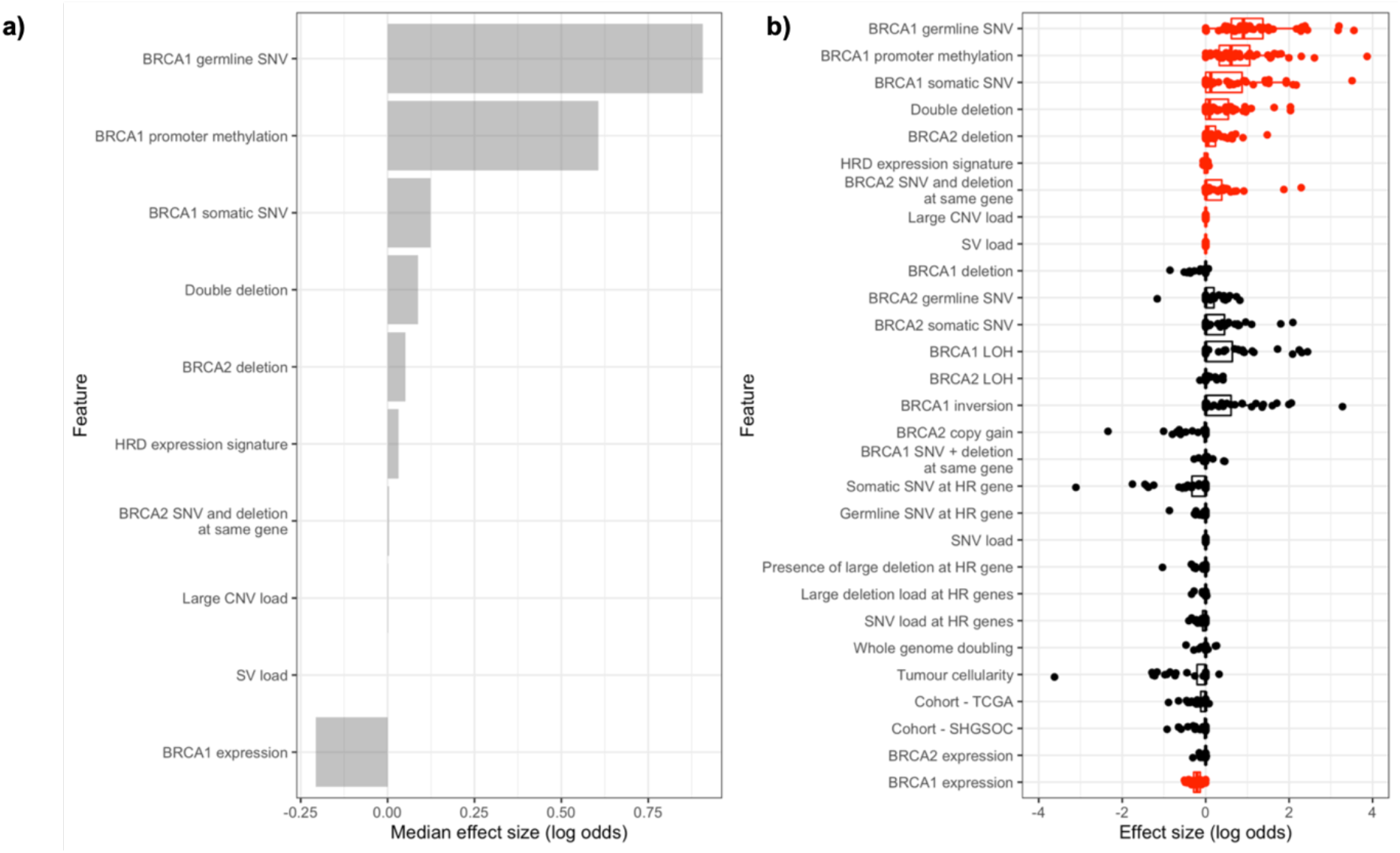
Integrative modelling of repair deficiency in HGSOC in full dataset. a) Median effect sizes of features selected to predict HRD, using elastic net regularised regression on 50 training/test set splits. Binary mutational status variables (e.g. presence/absence of BRCA1 somatic SNV) were included as factors and continuous variables were standardised to allow comparisons between variables. b) Distributions of effect size for each variable on HRD (log odds) in each training/test set split. Variables in red are selected for inclusion by the model in more than half of the training sets. It should be noted that, due to the lower number of samples with expression information and the increased number of features this model is likely to be underpowered to accurately identify significant features.

**Supplementary Figure 6.**
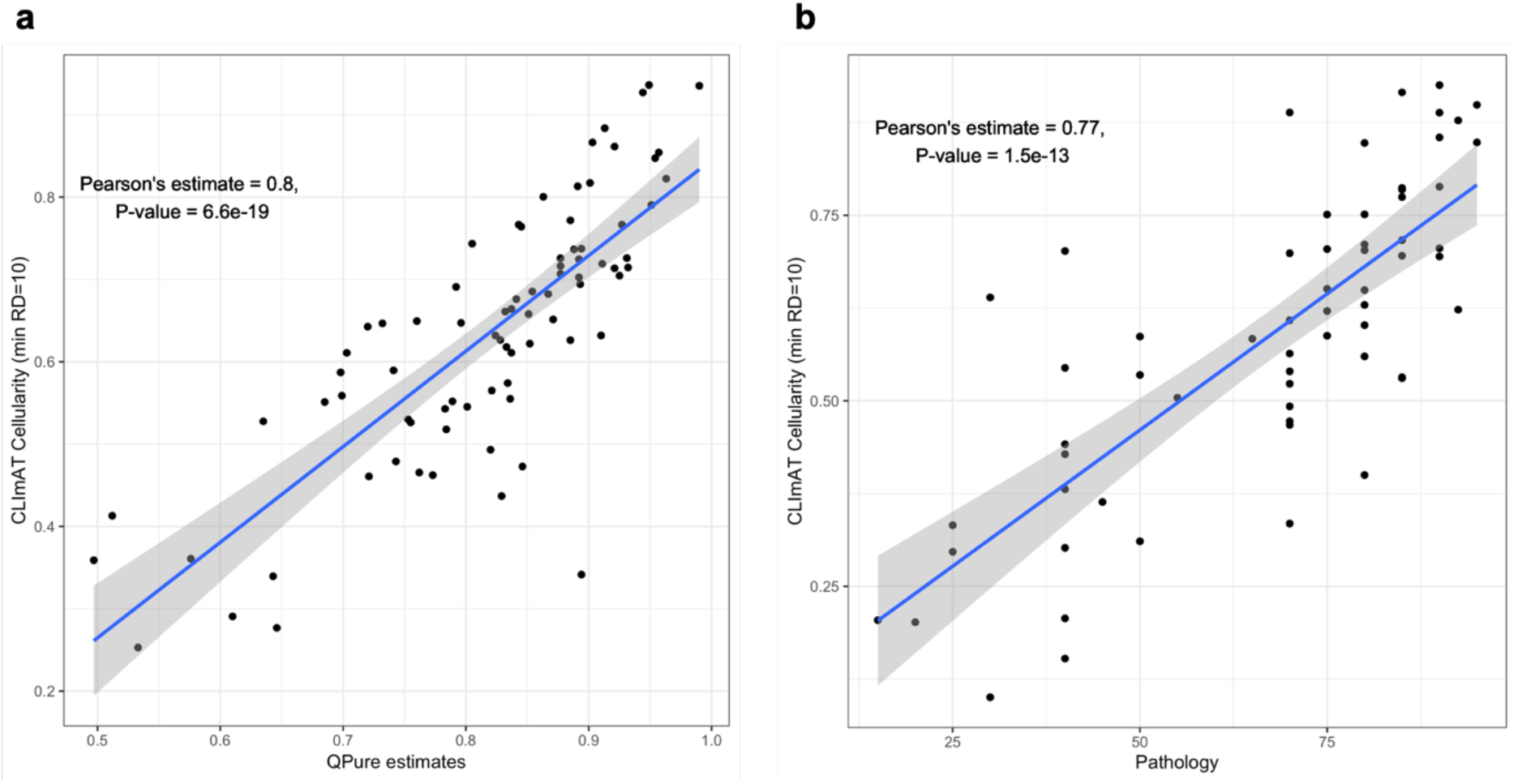
Comparison of methods to estimate the tumour cellularity in two cohorts. a) Estimates of tumour cellularity from allele-specific copy number tool CLImAT in comparison to estimates using qPure for the AOCS cohort. b) Estimates of tumour cellularity from allele-specific copy number tool CLImAT in comparison to scores from manual examination of the histopathology for the SHGSOC cohort.

